# Hyperdirect connectivity of opercular speech network to the subthalamic nucleus

**DOI:** 10.1101/2021.07.02.450909

**Authors:** Ahmed Jorge, Witold J. Lipski, Dengyu Wang, Donald J. Crammond, Robert S. Turner, R. Mark Richardson

**Affiliations:** Department of Neurological Surgery, University of Pittsburgh School of Medicine, PA 15213; Tsinghua University School of Medicine; Department of Neurobiology, University of Pittsburgh School of Medicine; Department of Neurosurgery, Massachusetts General Hospital, Boston, MA 02114; Harvard Medical School, Boston, MA 02115

**Keywords:** hyperdirect pathway, subthalamic nucleus, speech perception, speech production, deep brain stimulation

## Abstract

The importance of the basal ganglia in modulating cognitive and motor behaviors is well known, yet how the basal ganglia participate in the uniquely human behavior of speech is poorly understood. The subthalamic nucleus (STN) is well positioned to facilitate two basal ganglia functions critical for speech: motor learning and gain modulation. Using a novel paradigm to study cortical-subcortical interactions during speech in patients undergoing awake DBS surgery, we found evidence for a left opercular hyperdirect pathway in humans by stimulating in the STN and examining antidromic evoked activity in the left temporal, parietal and frontal opercular cortex. These high resolution cortical and subcortical mapping data provided evidence for hyperdirect connectivity between Broca’s area (typically corresponding to pars triangularis and pars opercularis of the inferior frontal gyrus) and the STN. In addition, we observed evoked potentials consistent with the presence of monosynaptic projections from areas of opercular speech cortex that are primarily sensory, including auditory cortex, to the STN. These connections may be unique to humans, evolving alongside the ability for speech.

**SIGNIFICANCE:** Using intracranial recordings from the basal ganglia and cortex in subjects undergoing deep brain stimulation, this study provides evidence for monosynaptic cortical inputs from motor planning, motor sensory, and auditory sensory cortices to the subthalamic nucleus. These observations suggest that in humans, the cortical-basal ganglia hyperdirect pathway is uniquely positioned to participate in speech production. Moreover, the existence of a monosynaptic connection between human sensory cortical areas and the subthalamic nucleus indicates a need to update traditional models of information transfer within cortical-basal ganglia-thalamocortical circuitry, which has significant implications for understanding other human cognitive behavior or dysfunction.

## INTRODUCTION

Possibly no human behavior requires more temporally precise control of multiple motor commands than speech. Speech neuroscience has traditionally focused on the cortex, but the importance of the basal ganglia in speech control is evidenced across an evolutionary scale. Mutations that affect basal ganglia development result in extreme deficits in speech motor-control and language comprehension (1), and damage to the adult basal ganglia can produce a variety of speech deficits (2). All regions of the basal ganglia share a common circuit plan, where the striatum receives topographically organized excitatory inputs from many cortical areas and conveys those inputs via direct and indirect pathways through the basal ganglia; this topography is largely conserved in outflow projections through the thalamus back to cortex. Thus, distinct motor, associative, and limbic functions are mediated via parallel cortical-basal ganglia-thalamocortical loops (3, 4). Among models of speech motor control, only state feedback control models(5), including the DIVA and GODIVA models (6), devote significant attention to function of basal ganglia-thalamocortical loops, yet no models of speech production consider the cortico-subthalamic hyperdirect pathway for cortical input to basal ganglia.

The hyperdirect pathway is a monosynaptic projection from the cortex to the STN (4, 7). Its presence was demonstrated recently in humans, by measuring evoked potentials (EPs) in sensorimotor cortex in response to low frequency STN stimulation (8). The recorded EPs include temporal components that group into three latency ranges: 1) very-short latency (< 2msec), consistent with transcortical motor evoked potentials (MEPs) in muscles, mediated by the corticospinal tract adjacency to the STN; 2) short-latency (2-10msec), consistent with antidromic activation of the hyperdirect pathway followed by cortico-cortical activation; 3) long latency (10-100msec), consistent with orthodromic cortical activation of basal ganglia-thalamocortical pathways. Although it has been hypothesized that output from the basal ganglia projects to Broca’s area (9) (disynaptically via thalamus), whether the basal ganglia receives direct input from Broca’s area remains uncertain. Based on Haynes and Haber’s demonstration of hyperdirect projections from multiple frontal regions in the primate (10), we hypothesized that the Broca’s area region, the left inferior frontal gyrus, has hyperdirect connection with the STN.

We recently described subthalamic nucleus (STN) single unit and local field potential activity during speech production (11, 12). Based on these studies and prior studies that demonstrated the functional connectivity of the STN to multiple cortical areas using simultaneous intracranial cortical-subthalamic recordings (13, 14), we hypothesized that cortical areas subserving not only speech production, but also perception, are connected to the STN via hyperdirect connections. Using electrocorticography (ECoG) in patients undergoing STN deep brain stimulation (DBS) implantation, we tested these hypotheses by recording cortical EPs to STN stimulation and mapping their location to stimulation sites within the STN. This method avoids the inherent limitations of diffusion-weighted magnetic resonance imaging approaches to tract tracing (15, 16). We found evidence for antidromic activation of premotor and motor regions of the frontal operculum. Remarkably, we found similar evidence for antidromic activation of sensory cortex, including auditory regions of the superior temporal gyrus.

Given that no tracing studies in nonhuman primates, to our knowledge, have described these pathways (17–20), hyperdirect connections of the cortical opercular speech network to the STN may be uniquely human.

## RESULTS

We successfully recorded and analyzed EPs from 20 patients with Parkinson’s disease, 17 with STN stimulation and 3 with GP stimulation. Cortical recording locations spanned multiple gyri and basal ganglia stimulation locations primarily spanned the sensorimotor territories of the STN or GP (Fig. 1 A-D).

**Fig. 1.**
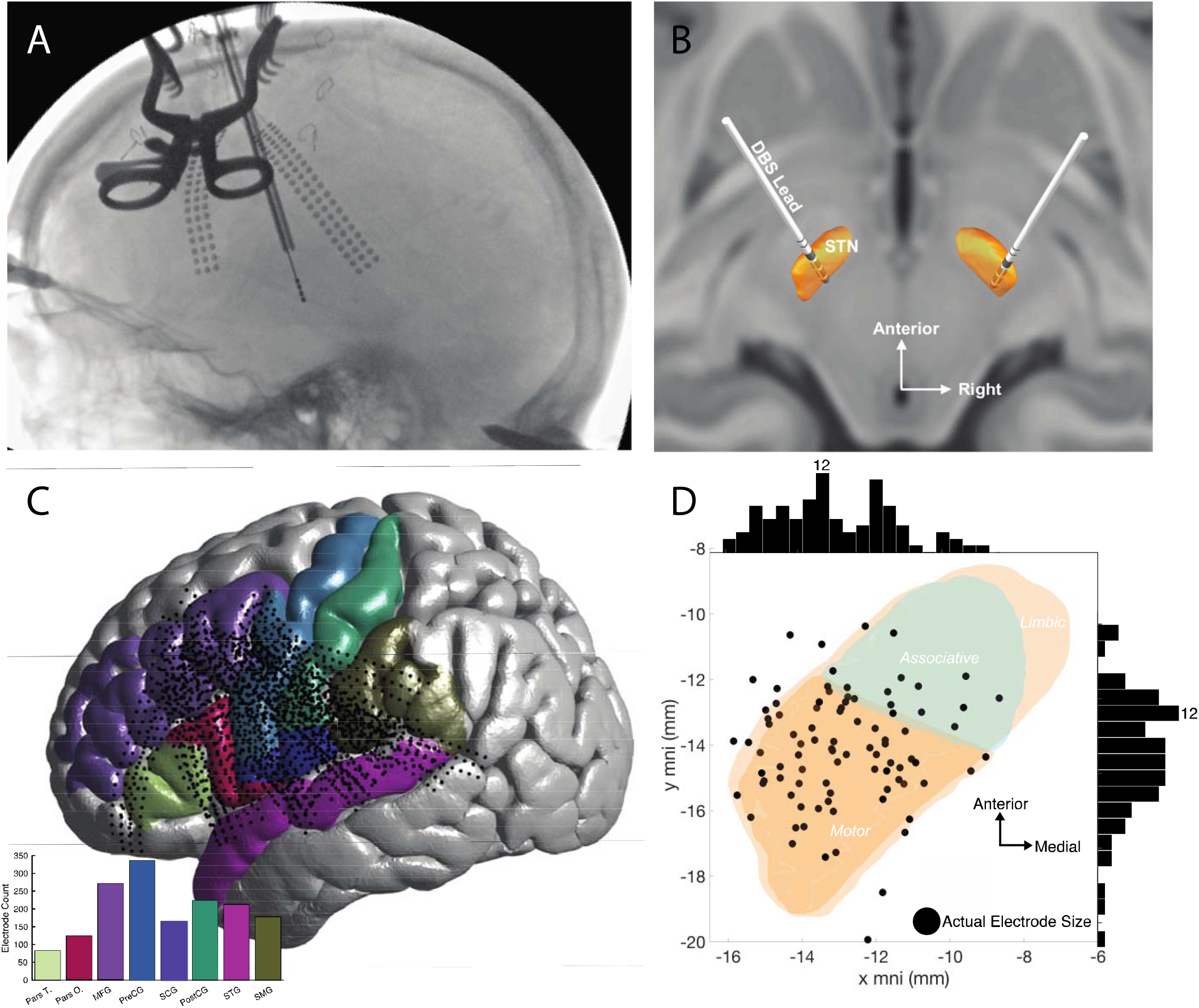
Experiment Setup. (A) Representative intraoperative fluoroscopy imaging performing during surgery to localize ECoG electrodes on the cortex. (B) Representative STN DBS localization. (C) Shows all ECoG electrodes locations (black circle dots) on the cortex with an inset histogram count of all electrodes per cortical area of interest (Destrieux atlas (25)). (D) All stimulation contact locations (black circle dots) within the left STN with sideline histograms highlight location distribution.

### Antidromic Cortical Excitation

We identified 9450 distinct voltage-time traces, each of which corresponded to a distinct combination of cortical ECoG recording location and STN stimulation location and amplitude. Three distinct short latency (2-10 ms) peaks related to antidromic activation were observed in each cortical region (Fig. 2 A), consistent with the existence of a fast-conducting monosynaptic connection. These peaks were present not only in premotor and motor regions (8, 28), but also in the SMG and STG. No distinct peaks were observed during GPe or GPi stimulation in the 2-10 second latency period. Long-latency EPs in the 20-100 ms latency period were consistently recorded to both STN and GP stimulation. These two long-latency peaks (see example in Supplementary Figure 1) did not change for different STN stimulation locations (peak latency for the two associated peaks was 39.5±0.7 ms and 60.1±4.2 ms, respectively); however, stimulation of the GPe resulted in slower peaks (peak latency for the two associated peaks is 24.1±0.2 and 46.3±2.6, respectively) than stimulation of GPi (peak latency for the two associated peaks is 17.6±0.1 and 38.8±2.4ms, respectively). consistent with orthodromic activation via the basal ganglia-thalamocortical loop. To quantify the average cortical evoked response in the 2-10 ms range from STN stimulation, we averaged all traces (n=9450) (Fig. 2 B). The average amplitude and latency responses were 1.8±2.9 μV and 3.1±0.8 ms for EP1, 1.5±2.7 μV and 4.9±1.3 ms for EP2, and 1.1±2.3 μV and 6.5±1.8 ms for EP3.

**Fig. 2.**
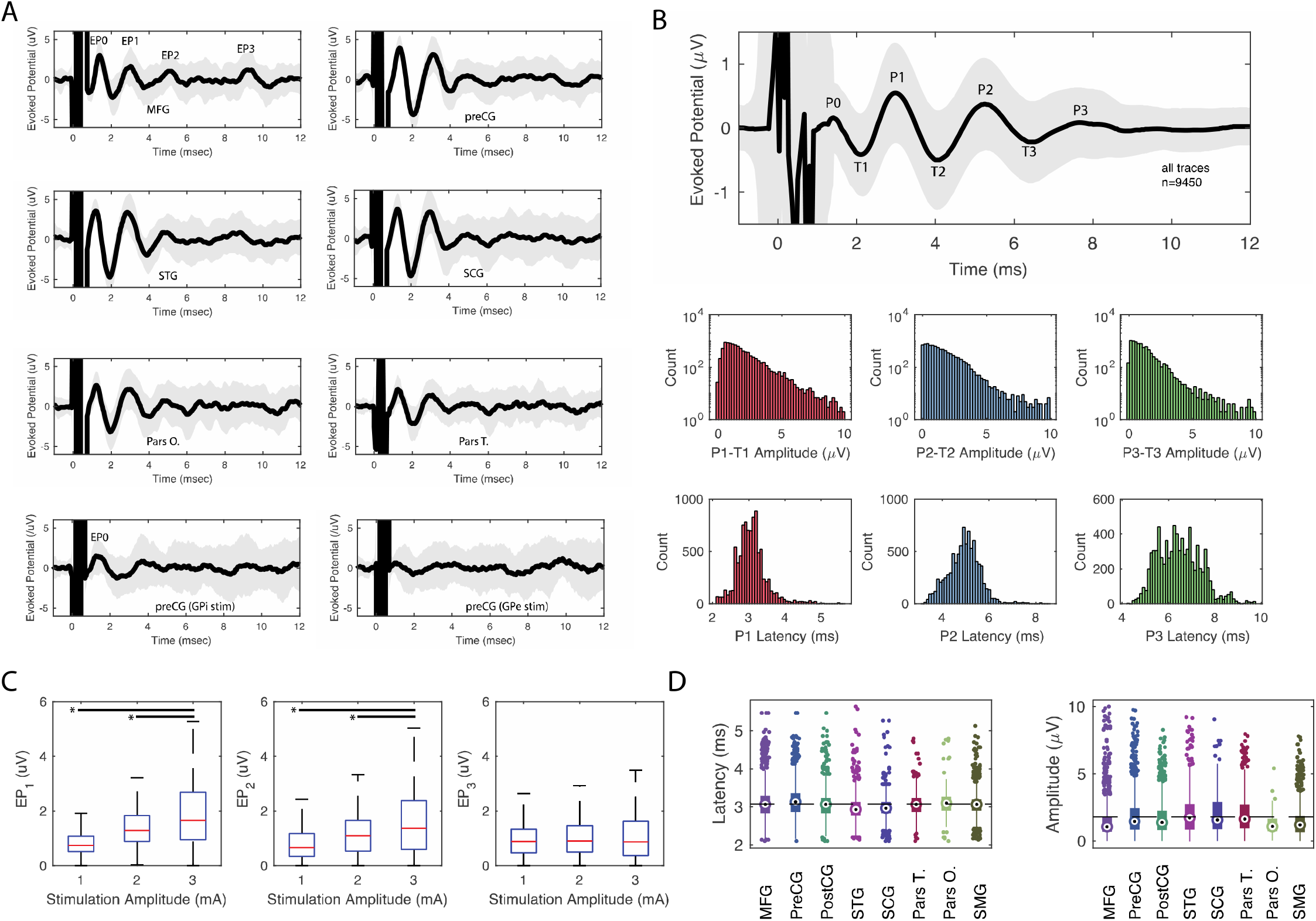
Characterization of cortical evoked potentials. (A) Examples of STN (top six subpanels) and GP (bottom two subpanels) stimulation, shown here at 3mA, in a single subject for different brain gyri. GPi stimulation elicited a very-short evoked potential (EP0) consistent with its proximity to the internal capsule but no appreciable EP1-3. GPe stimulation elicited no EPs. Thick black line and grey region corresponds to the mean and standard deviation of the cortical evoked potential of 30 stimulation trials for one subject and one stimulation location. Stimulation occurred at time 0. EP0 corresponds to the peak associated with very short evoked potentials while EP1-3 corresponds to the peaks involved with the hyperdirect pathway. (B) Averaged EP peaks (P) and throughs (T) after STN stimulation for all traces, mean as a black line, standard deviation as a gray shadow. (C) Amplitude and latency distribution of the difference between peaks and throughs, or EP 1-3. (D) Evoked potential amplitude (with interquartile ranges) changes are plotted as a function of stimulation amplitude for different evoked potential peaks. *=significant difference in an ANOVA test. (E) Cortical evoked potential (EP1) latency and amplitude distribution plotted as a function of cortical area (I) latency and (J) amplitude for all traces. Abbreviations: MFG: middle frontal gyrus, preCG: precentral gyrus, postCG: postcentral gyrus, STG: superior temporal gyrus, SCG: subcentral gyrus, Pars T.: pars triangularis, Pars O.: pars opercularis, SMG: supramarginal gyrus. On panel (D), Circle = median, large rectangle represents 25-75 quantiles, thin line represents upper and lower quartiles, respectively.

To quantify the STN stimulation threshold (and resulting EP peak magnitude), we tested the effects of different stimulation amplitudes on EP amplitude (Fig. 2 C) in different cortical locations (Supplementary Figure 2). EP1 and EP2 amplitudes were dependent on stimulation intensity up to 3.0 mA (F_2,9085_=1408, p<0.001 and F_2,8650_=505, p<0.001, ANOVA, respectively) while the EP3 did not show variation with different stimulation amplitudes (F_2,7934_=4.5, p=0.11, ANOVA). We did not test stimulation with currents higher than 3mA, given that the average EP standard deviation with 3mA stimulation was 2.84 μV. Stimulations of 1.0 and 2.0 mA were shown to be sub-threshold given that stimulation at 3.0 mA produced significantly higher EP amplitudes. Moreover, given that the average EP standard deviation with 3mA stimulation was 2.84 μV, we decided to perform all subsequent analysis at a stimulation intensity of 3mA. The reason to not perform stimulations at a higher amplitude than 3.0 mA was to maintain recordings of robust cortical EP responses (i.e. higher than sub-threshold) while minimizing voltage spread within the STN. In addition, as the neural basis of the circuitry which generated the long-latency EP’s can only be speculated, we focused the analysis entirely on the EP1 component as it is likely the only evoked potential component arising from activation of the hyperdirect pathway alone (8, 28–30).

Next, we investigated whether the wide range of EP1 latencies and amplitudes varied as a function of cortical location. We found that EP1 amplitude varied significantly between cortical locations (F_7,6146_=210, p-value<0.001, ANOVA) with voltage mean and standard deviation for preCG (1.9±1.5μV), MFG (1.5±1.5μV), postCG (1.8±1.4μV), pars T. (1.2±0.7μV), pars O. (2.1±1.6 μV), STG (2.0±1.5μV), SCG (2.0±1.4 μV), and SMG (1.6±1.3μV) gyri shown in Fig. 2 D. These voltage differences suggested that different locations within the STN receive a different proportion of efferent axons from different cortical areas. A similar analysis for latency did not reveal significant differences. The stimulation voltage variance for each area exhibited a broad range of voltage amplitudes, which suggests that the nature of the hyperdirect connection also depends on the precise location within the STN.

### Cortical evoked potential amplitude depends on STN stimulation location

We observed wide EP ranges for each cortical area (Fig. 2 D). To better understand this variability, we separated each EP as a function of cortical area and STN stimulation location. We found that for the precentral gyrus, the EP amplitude was a function of STN stimulation location (Fig. 3 A-C). For the precentral gyrus cortical EPs, STN stimulation locations closer to the center of the STN motor region, as defined by the DISTAL subcortical atlas, produced a significantly higher EP voltage than when stimulating closer to the center of the STN associative region (Spearman’s rho=−0.53, p<0.001). Contrastingly, STN stimulation closer to the STN associative region center produced higher EPs in the middle frontal gyrus (Fig. 3, D-F) when compared to stimulations closer to the STN motor region (Spearman’s rho=0.41, p=0.003). The estimated distance between these two centers is 2.1 mm.

**Fig. 3.**
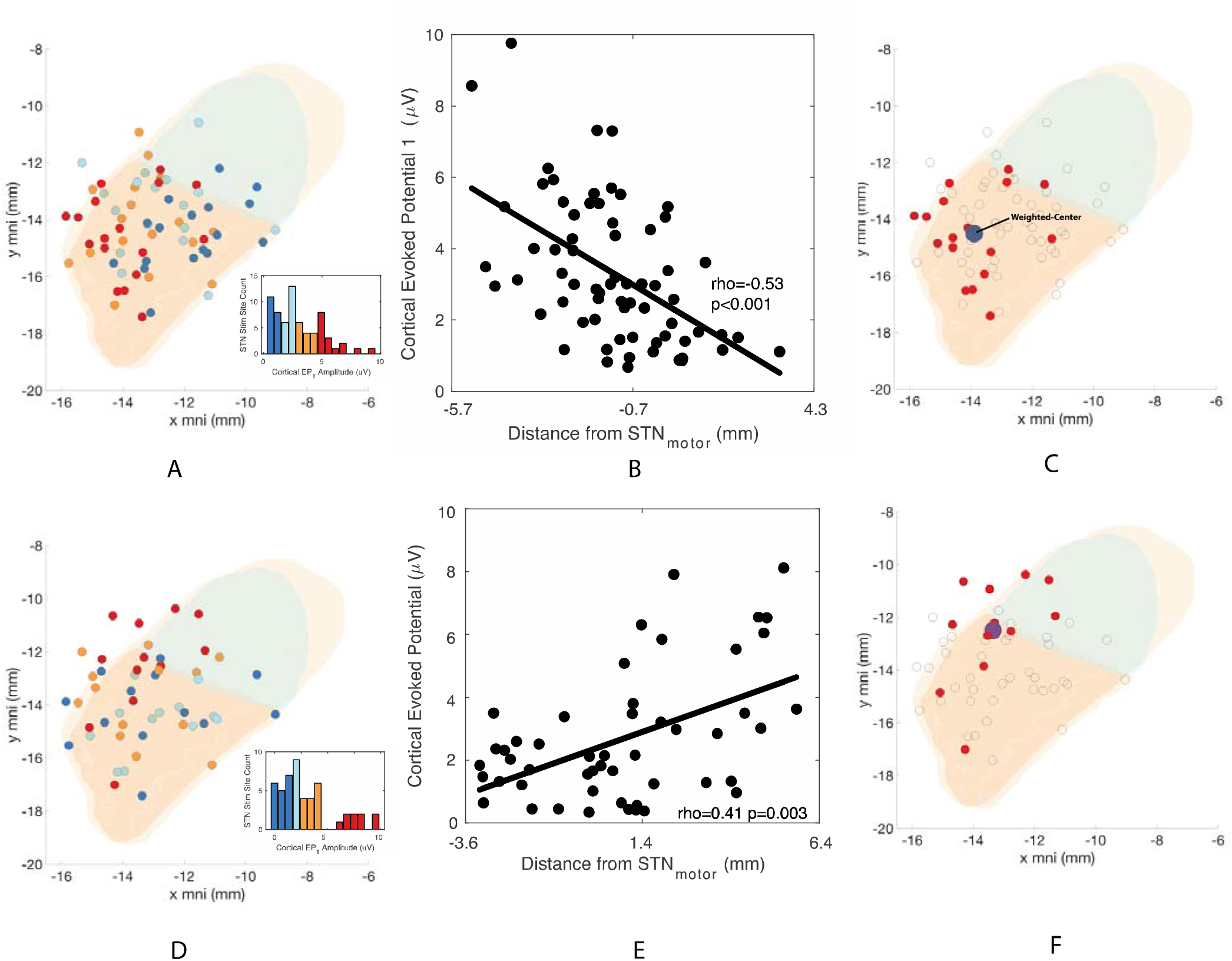
Cortical evoked potentials (EP) projected onto the STN. (A) Individual STN stimulation site and projected average precentral gyrus cortical EP in color representing interquartile voltage measures (dark blue, 1^st^ quantile, light blue, 2^nd^ quantile, orange, 3^rd^ quantile and red, 4^th^ quantile. (B) All precentral gyrus cortical evoked potential projected onto the defined one-dimensional STN axis (C) Weighted center of evoked potential response for the upper quartile of precentral cortical evoked potentials. (D-F) are similar in definition as previous figures but projecting the cortical EP from the middle frontal gyrus. The motor aspect of the STN is colored in orange, associative in green and limbic in yellow in panels A,C,D,F.

By visualizing EP1 on the cortex, we found that stimulations closer to the center of the STN motor region produced higher EP1 in cortical regions closer to the central sulcus (Fig. 4 A); while stimulations closer to the center of the STN associative region produced higher EP1 in cortical regions farther away from the central sulcus (Fig. 4 B). In order to summarize the overall findings for STN sub-regions receiving input via the hyperdirect pathway, we plotted the weighted center of the average cortical evoked potential amplitude (from Fig 3 A-F) for each cortical area onto the STN map and projected them onto the STN axis (Fig. 4 C-F). We took this STN axis location and compared it to the distance from the central sulcus (a fronto-posterior simplification of the cortex). We found that there was a significant relationship between STN location (motor to associative region) and distance from central sulcus (Spearman’s rho=0.80, p=0.01). In other words, stimulation of the STN motor region produced the highest voltages in cortical regions closer to the central sulcus (e.g. preCG, postCG and SCG) which strongly suggests that these cortical areas project the highest density of efferent fibres directly to innervate STN, while stimulation of STN associate regions produced the highest voltages in cortical regions further away from the central sulcus (e.g. MFG, Pars T. and SMG).

**Fig. 4.**
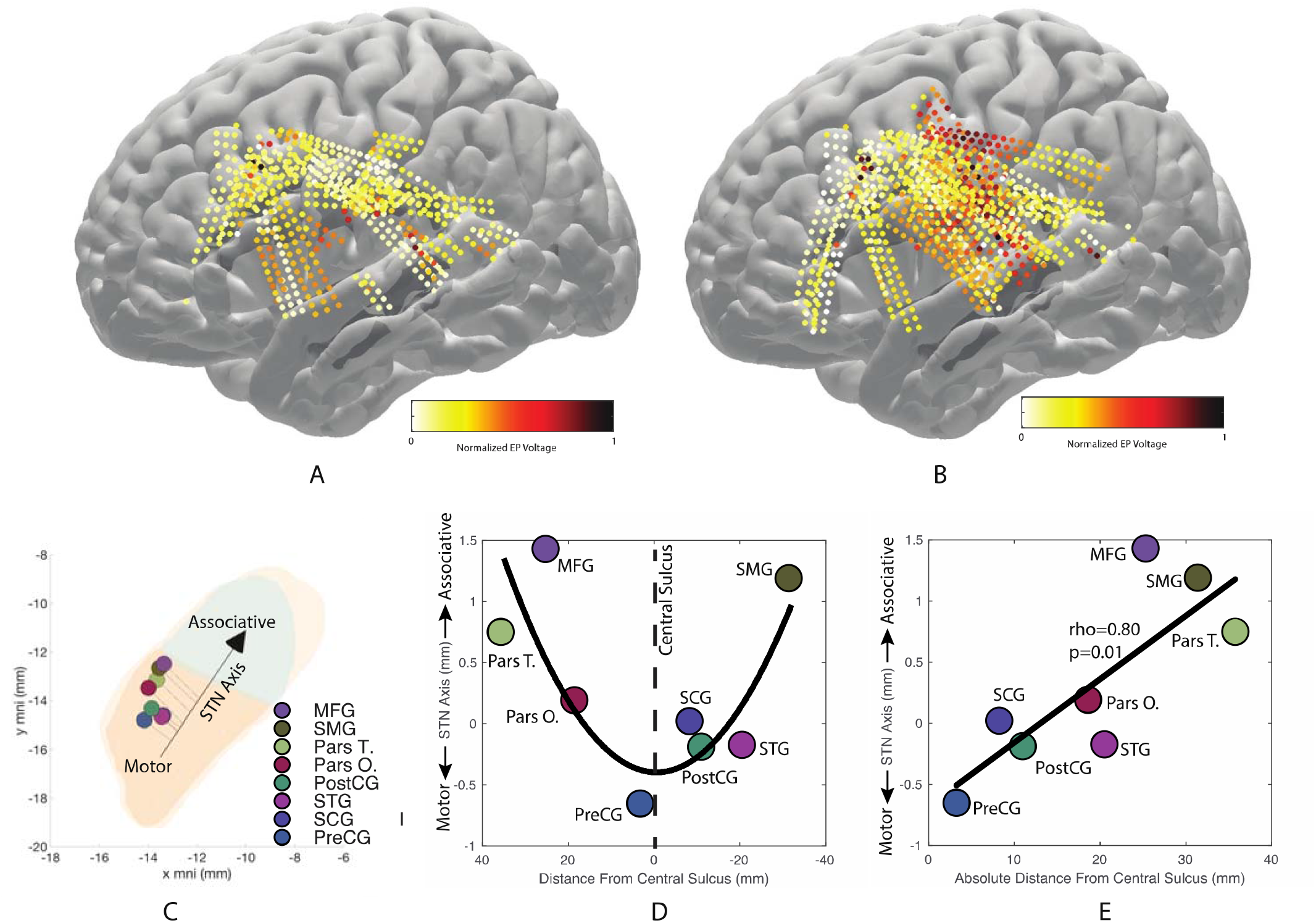
Evoke Potential Amplitude as a Function of STN stimulation location. (A) EP amplitude for STN stimulations closer to the STN associative center (the quantile containing the most frontomedial stimulation locations in the STN). (B) EP amplitude for STN stimulations closer to the STN motor center (the quantile containing the most posterolateral stimulation locations in the STN). Electrodes that did not show an EP are not shown. (C) Weighted-center of cortical evoked potential in the STN per cortical area; motor aspect of the STN colored in orange, associative in green and limbic in yellow. (D) Quadratic relationship between the STN center-of-voltage and the corresponding cortical location on the y mni axis. (E) Model depicting a significant linear relationship between STN center-of-voltage and the absolute distance from the central sulcus. Legend: (Pars O.), opercular part of the inferior frontal gyrus, (Pars T.), the triangular part of the inferior frontal gyrus, (MFG), the middle frontal, (PreCG), precentral, (PostCG), postcentral, (SCG), subcentral, (SMG), supramarginal and (STG), superior temporal gyrus.

## DISCUSSION

We used high-density electrocorticography to record directly from the cortical surface in patients undergoing DBS implantation to demonstrate that all areas of the cortical opercular speech network, including sensory areas, are monosynaptically connected to the STN. We confirmed physiological evidence of hyperdirect (monosynaptic) connectivity between the IFG and STN in humans, recently demonstrated by Starr and colleagues(28). Importantly, we extended those findings of antidromic activation of motor and prefrontal cortex to include sensory cortex (post-central gyrus), specifically demonstrating the occurrence of evoked potentials that indicate monosynaptic connections between sensory cortical areas involved in speech processing and the STN. In agreement with the idea that input to the STN is segregated within topographically organized loops, with a degree of overlap, we found that stimulation closer to the central motor territory of the STN was more likely to produce EPs in cortical areas closer to the central sulcus, while stimulation closer to the associative territory of the STN was more likely to produce EPs in cortical areas further from the central sulcus.

### Evidence for Hyperdirect Pathway Projections to the Opercular Cortical Speech Network

Given that non-human primates do not speak, and that viral synaptic tracing studies cannot be performed in humans, EP mapping during STN DBS surgery is the only method that can directly characterize a speech cortex to STN hyperdirect pathway in humans. Mapping the hyperdirect pathway from speech-related cortical areas is important for several reasons: 1) these areas enable behavior that is uniquely human, and thus their connectivity to subcortical nuclei may be uniquely human. 2) the hyperdirect pathway has been proposed to administer a brake signal by stimulating GPi, yet it also may administer a GO signal by stimulating GPe, indirectly inhibiting GPi; understanding the origin of direct inputs to the STN can inform hypothesis generation regarding the functional roles of these inputs(31, 32). 3) hyperdirect connectivity likely signifies the presence of loops that allow information in speech-specific cortex to be transferred directly to the direct and indirect basal ganglia pathways, in order to modulate speech production.

We showed the occurrence of three clusters of STN stimulation-evoked cortical potential peaks in the 2-10 ms range, previously described in EEG (29, 30, 33, 34), ECoG (8, 35), MEG (36) and finite-element modeling (37) studies as consistent with antidromic cortical activation via the hyperdirect pathway. Indeed, the latencies of these peaks were too fast (2-10 ms) to be of orthodromic origin via the globus pallidus and thalamus (a relay that has a duration on the order of 10-40 ms) and too slow to be related to corticospinal pathways (<1.5 ms) (8, 30). We focused on the first EP peak since it is putatively the most significant peak associated with the hyperdirect pathway. Although the second and third EP peaks observed could be associated with slower and less myelinated hyperdirect pathway fibers, these peaks could also be the product of cortico-cortical interactions after the first antidromic impulse, a multiphasic response from an activated GP-STN pathway (38) or the product of an STN-to-cortex direct pathway (39).

We showed that these potentials can be evoked from STN across broad areas of cortex. Although the majority of afferents to the STN come from the GPe, it has been argued that the hyperdirect pathway can carry key inputs to alter motor and non-motor functions mediated by the basal ganglia (38, 40–42). In addition, computational models have shown that DBS stimulation can robustly propagate signals to the motor cortex with high fidelity (37). Here we show involvement of other cortical areas, particularly areas involved with speech production. In agreement with our results, the left inferior frontal cortex (including pars triangularis and pars opercularis), hypothesized to part of the basal ganglia thalamocortical circuitry (43), has been suggested to be directly connected to the STN by human tractography and fMRI studies (44).

### STN Hyperdirect Pathway is Topographically Organized

The basal ganglia topographical organization with motor, associative and limbic circuitry having distinct regions across the putamen, GPe and GPi has been well described (3, 45, 46). Here we show that ventromedial STN stimulation produces higher EPs in the premotor cortex while dorsolateral STN stimulation produces higher EPs in the motor cortex, consistent with anterograde tracer studies in nonhuman primates (10). Moreover, there is a relationship between STN stimulation location and the estimated cortical EP distance from the central sulcus, suggesting a precise topography of cortical innervation of the STN. In addition to shedding light on STN function, hyperdirect pathway topology is also of clinical importance, given that stimulation of different STN sub-regions has been associated with non-motor behavioral responses, in patients with PD (47, 48).

Rapid modulation of the control of speech requires highly coordinated information transfer between planning, production and perception hubs of the speech network. Hyperdirect connectivity of premotor, motor-sensory, and auditory cortical regions suggests that the basal ganglia may play an integral role in state feedback control of speech production, where auditory information is compared with a prediction derived from efference copy of motor output (5). The basal ganglia are well suited to modulate these processes (6, 49), and have been implicated in processing of prediction error(50–52), which in the context of speech production would contribute to an internal model tracking the state of the vocal tract. In addition to a potential role in evaluating efference copy, the STN could participate in speech gain modulation (12, 53), via either movement inhibition or a prokinetic effect (54). The relay of sensory information directly to the STN could help shape this modulation. Indeed, we have previously shown evidence for indirect information flow between primary sensory cortex and the STN during hand gripping movement (13, 55). The existence of a sensory hyperdirect pathway, however, has not previously been hypothesized. Studies in rat models (38, 56) stimulating the cortex and measuring STN spikes and LFPs have shown evidence that there is a direct connection between prefrontal, premotor, cingulate, M1 and S1 but a similar response was not observed with STG stimulation. In contrast, a tractography study in humans did suggest the presence of a hyperdirect connection from prefrontal, M1, S1 and STG to the STN (57). It is possible, therefore, that hyperdirect connections to STN from sensory cortical areas, ultimately involved in processes related to speech perception, have co-evolved with speech ability in humans.

### Limitations

We tested stimulation amplitudes in the 1-3mA range (see Supplemental Figure 2), since the variance of the evoked potential at 3mA became large. We did not reach an asymptote supratreshold, and thus we cannot be certain that 100% of the hyperdirect efferents were activated. We understand that the EP amplitude represents the amplitude of the net dipole summed over thousands of local dipoles in the region covered by our ECoG grid. We assumed that all local dipoles were sampled evenly across all the available cortical coverage as shown in Fig. 1; we are not taking into consideration dipoles occurring outside the ECoG coverage area or dipoles in the sulci region (due to an inherent lack of ECoG coverage).

## CONCLUSION

Using intracranial recordings from the basal ganglia and cortex in subjects undergoing deep brain stimulation, this study expands the known monosynaptic cortical inputs to the human subthalamic nucleus to include the entire frontal-parietal-temporal opercular cortex, including auditory cortex. These data provide evidence for a hyperdirect pathways from motor planning, motor sensory, and auditory sensory cortices to the basal ganglia that are uniquely positioned to participate in speech production.

## METHODS

### Participants and Surgery

Twenty patients undergoing STN (n=17), globus pallidus internus (GPi, n=3) DBS surgery were enrolled in an IRB-approved protocol, following informed consent. All patients were right-handed with presumed left language dominance. Dopaminergic medications were held for 12 hours prior to surgery. The STN or GPi was targeted utilizing standard clinical techniques in awake, microelectrode-guided surgery (21, 22). It is the surgeon’s practice to always begin with the left side first, and all data was collected in the left hemisphere. Prior to insertion of microelectrodes, one or two subdural high-density ECoG arrays (63 channels, 1 mm contact diameter, 3mm separation, 3×21 contact configuration; PMT, Chanhassen, MN, USA) were temporarily placed at prespecified cortical locations through the single standard burr hole. The ECoG Strip location was planned preoperatively to span cortical areas including the frontal operculum and/or the premotor, ventral sensorimotor, and superior temporal cortex.

### STN/GPi Stimulation and ECoG Recordings

Clinical microelectrode recording was completed using three simultaneously placed Alpha Omega Microprobe electrodes (Alpha Omega Co, Alpharetta, GA, USA) in the center, medial and posterior trajectories of the Ben-Gun array. STN or GPi monopolar stimulation was conducted for clinical mapping purposes through the macro cylindrical contact (“ring electrode”, diameter 0.7mm, length 1mm) using the Neuro Omega (Alpha Omega Co, Alpharetta, GA, USA) stimulation software. Upon completion of clinical testing, research stimulation was conducted at 1Hz for 30s (30 stimulation pulses) or 10 Hz for 30 s (300 stimulation pulses) at stimulation intensities of 1, 2 and 3 mA, at two separate depths within the STN, separated by at least 2 mm in the z-dimension. Simultaneous with stimulation, cortical evoked potentials were recorded, amplified, and digitized using a Grapevine Neural Interface Processor (Ripple Neuro, Salt Lake City, UT, USA). Signals were recorded at sampling rate of 10 kHz with all channels referenced to scalp ground.

### Localization of ECoG recordings and STN stimulation leads

ECoG strip location was determined with intraoperative fluoroscopy imaging or CT imaging, preoperative in-frame CT, and preoperative MRI as previously described (23). Normalization of each subject’s MRI to MNI brain space and automatic identification of ECoG electrode and gyri location was conducted a posteriori with FreeSurfer and the Destrieux atlas (24, 25). According to this atlas, eight anatomical categories were covered by ECoG recordings in this study, including the opercular part of the inferior frontal gyrus (Pars O.), the triangular part of the inferior frontal gyrus (Pars T.), the middle frontal (MFG), precentral (PreCG), postcentral (PostCG), subcentral (SC), supramarginal (SM) and superior temporal gyrus (STG). STN contact locations were determined using Lead-DBS MNI ICBM atlas and software (26). DICOM images were coregistered using Advanced Normalization Tools (ANTs) and hybrid statistical parametric mapping (SPM) algorithms and then normalized to the International Consortium for Brain Mapping (ICMB) 152 nonlinear 2009b template. Coordinates were recorded in MNI space and STN geometry (i.e. motor, associative, limbic) was defined by the DISTAL atlas (27).

### Signal Preprocessing

Stimulation start times were assigned by identifying the first time-bin with largest voltage deflection on a channel displaying a large stimulation artifact. The remaining ECoG channels were aligned to these stimulation times and a trial was defined by each stimulation time. To filter out low frequency fluctuations without introducing filter artifacts, raw voltage values for each trial were de-trended by subtracting a low-order polynomial fit of the signal (8^th^ order). For 1 Hz stimulation, 30 trials within each session were averaged per channel and then smoothed using a 5-bin (0.17ms) moving window (8). The first positive voltage peak deflection (peak 0) after stimulation was defined as cortical evoked potential 0 (EP0), with subsequent voltage peak deflections labeled as EP1 through EP3. Each temporal component of the evoked potential was separated into the peak and trough of each response accordingly; for example, cortical evoked potential 1 (EP1) amplitude and latency were defined as the amplitude from trough 1 (T1) to peak 1 (P1) and the latency was defined at the peak of P1. All data processing and analysis was performed using custom code in MATLAB 2017 (Mathworks, Natick, MA).

### Weighted-Center of Cortical Evoked Potential Calculation

To determine the location in the STN that receives the strongest hyperdirect projection from each cortical area, a weighted-center of voltage calculation was taken. This calculation is analogous to a center of mass calculation, with voltage (in our case, EP1) being the weight for each STN stimulation location. In this case, **R** is defined as the vector that points to the center of weighted EP. In other words, **R** is the location at which the STN would produce an average cortical response in the cortical area of interest based on the sum of all actual stimulation locations, **r**_i_, multiplied by the recorded evoked cortical responses, v_i_.

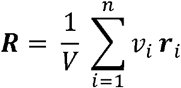

To simplify the dimensionality of the STN space, we defined a one-dimensional STN axis with two points, namely the center of the STN associative region volume (as described in [name] atlas) (MNI coordinates = [−10.4 −11.7 −7.6] mm) and center of STN motor region volume (MNI coordinates = [−12.6 −15.0 −7.1] mm) with origin at the STN center.

### Statistical Analysis

Analysis of variance (ANOVA) was performed with multiple comparison correction using the Tukey-Kramer method.

## ACKNOWLEDGEMENTS

Funded by NINDS U01NS098969 (WJL, DJC, RST, RMR)

## SUPPLEMENTARY MATERIALS

**Supplementary Figure 1.**
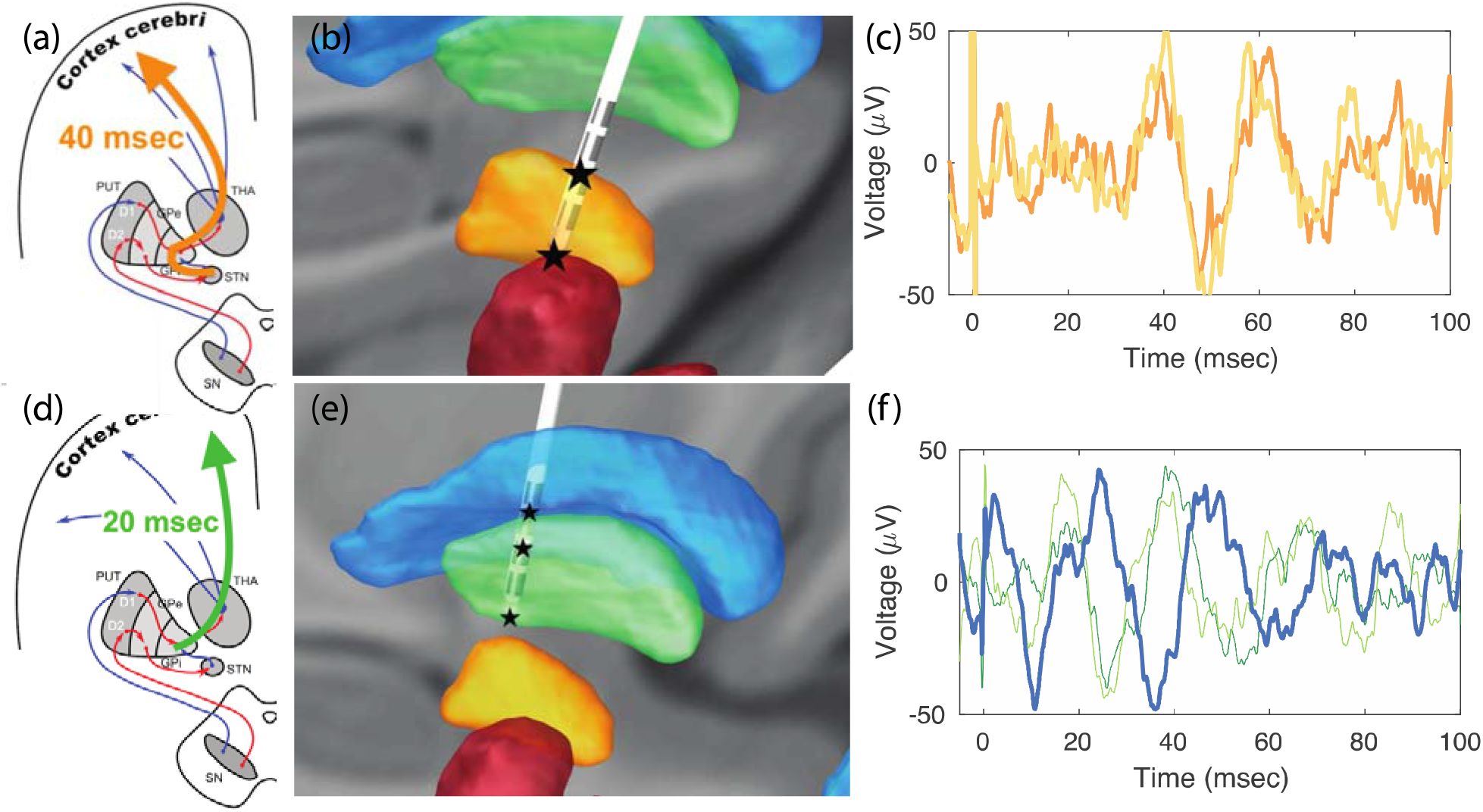
Long latency evoked potentials in (a-c) STN and (d-f) GPi. These long latency evoked potentials are consistent with the orthodromic pathway.

**Supplementary Figure 2.**
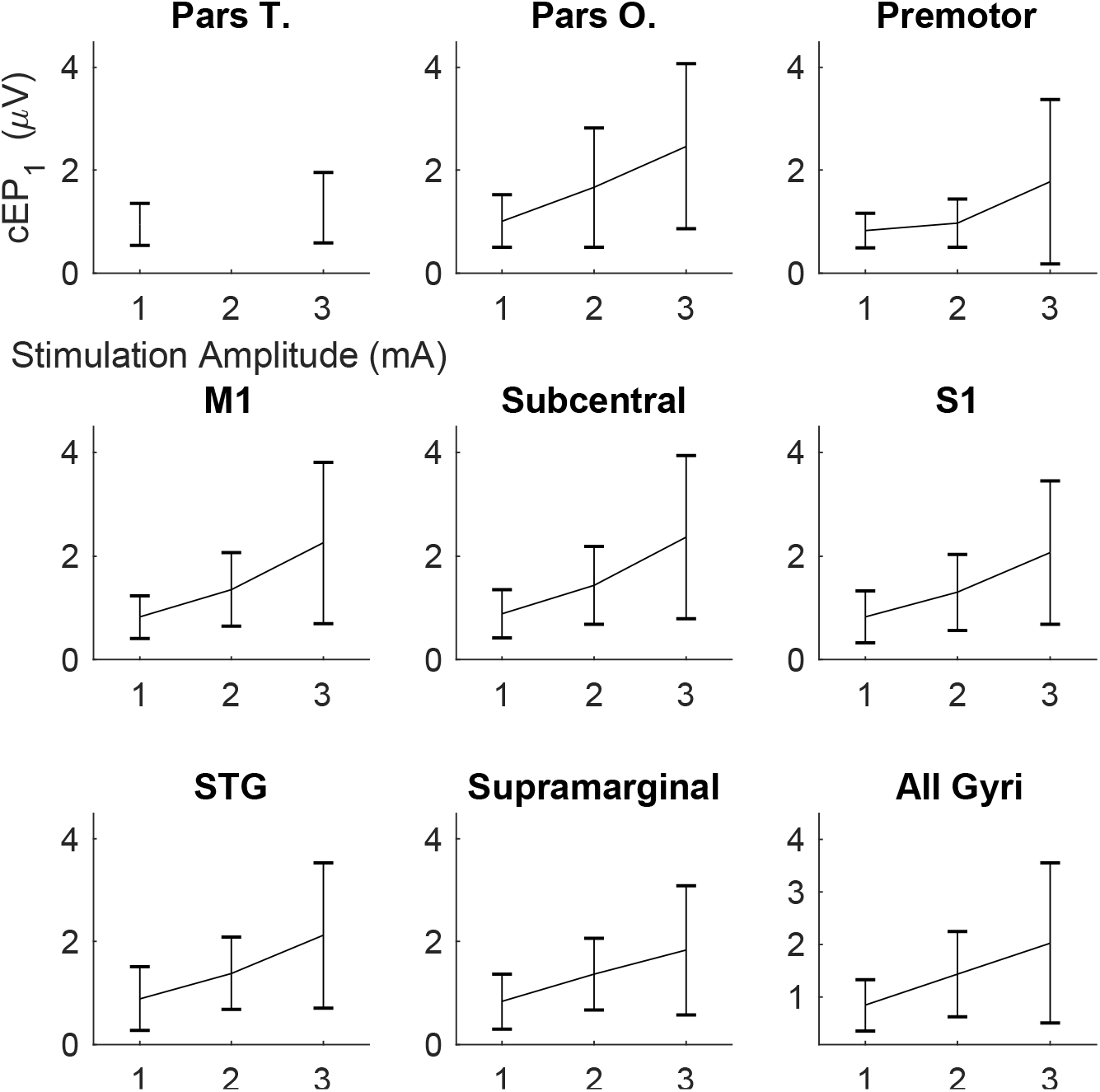
Evoked potential amplitude (+/− standard deviation) changes as a function of stimulation amplitude for different cortical areas.

